# Colchicine inhibited oxidative stress-induced endothelial cell senescence via blocking NF-κB and MAPKs: Implications in vascular diseases

**DOI:** 10.1101/2023.08.04.552075

**Authors:** Huakang Zhou, Dilaware Khan, Sajid Muhammad Hussain, Norbert Gerdes, Carsten Hagenbeck, Majeed Rana, Jan Frederick Cornelius, Sajjad Muhammad

**Affiliations:** Department of Neurosurgery, Medical Faculty and University Hospital Düsseldorf, Heinrich-Heine-Universität Düsseldorf, Germany; Cologne Center for Genomics (CCG), University of Cologne, Weyertal 115b, 50931 Cologne, Germany; Division of Cardiology, Pulmonology and Vascular Medicine, Medical Faculty and University Hospital, Heinrich-Heine-University Düsseldorf, Germany; Cardiovascular Reasearch Institute Düsseldorf (CARID), Medical Faculty and, Heinrich-Heine-University Düsseldorf, Germany; Clinic for Gynecology and Obstetrics, University Clinic, 40225 Düsseldorf, Germany; Department of Oral-, Maxillofacial and Facial Plastic Surgery, University Hospital Düsseldorf, Moorenstrasse 5, 40225 Düsseldorf, Germany; Department of Neurosurgery, University Hospital Helsinki, Topeliuksenkatu 5, 00260 Helsinki, Finland; Department of Neurosurgery, King Edward Medical University, Lahore, Pakistan

**Keywords:** Oxidative stress, endothelial cells, cellular senescence, inflammation, colchicine, NFκ-B, MAPKs, mTOR

## Abstract

Smoking, alcohol abuse, and hypertension are – among others the potential risk factors for cardiovascular diseases. These risk factors generate oxidative stress and cause oxidative stress-induced DNA damage, resulting in cellular senescence and senescence-associated secretory phenotype (SASP). The SASP factors in feed-forward response exacerbate inflammation and cause tissue remodeling, resulting in atherosclerotic plaque formation and rupture. Colchicine was used to ameliorate oxidative stress-induced senescence in human umbilical vein endothelial cells. Oxidative stress was quantified by reactive oxygen species (ROS) assay and oxidative stress-induced DNA damage was analyzed by 8-OHDG immunofluorescence staining. Endothelial cell senescence was visualized by β-gal staining. The relative mRNA expression was quantified by qPCR and protein analysis was performed by Western blot. Colchicine inhibited ROS generation and mitigated oxidative stress-induced DNA damage. It dampened oxidative stress-induced endothelial cell senescence and improved the expression of DNA repair protein KU80 and aging marker Lamin B1. The drug attenuated the expression of senescence marker P21 at mRNA and protein levels. The pathway analysis showed that colchicine inhibited NF-κB and MAPKs pathways and subdued mTOR activation. Colchicine also attenuated mRNA expression of interleukin (IL)-1β, IL-6, IL-8 MCP-1, ICAM-1, and E-selectin. Furthermore, colchicine reduced the mRNA and protein expression of matrix metalloproteinase (MMP-2). In summary, colchicine blocked oxidative stress-induced senescence and SASP by inhibiting the activation of NF-κB and MAPKs pathways.

## Introduction

State-of-the-art in cardiovascular research suggests that cellular senescence contributes to the initiation and progression of cardiovascular diseases including atherosclerosis, aneurysms, arterial stiffness, hypertension, and heart failure ^1^. The cells can divide into a finite number after which they become replicative senescent ^2, 3^. These cells stop dividing and reach a state of growth arrest, called the Hayflick limitation ^3^. In addition to this replicative senescence, other stimuli including acquired risk factors for cardiovascular diseases such as alcohol abuse and smoking can also induce premature senescence in endothelial ^4, 5^ and other cell types. The phagocytic activity of monocytes and neutrophils is impaired with increasing age, resulting in the accumulation of senescent cells in the vascular tissue. This accumulation of senescent cells can negatively affect life span and promote age-related diseases including cardiovascular diseases ^1, 6, 7^. The elimination of senescent cells improved health- and life-span and protected against cardiovascular diseases ^6, 7^. Previous studies have shown the accumulation of senescent cells in atherosclerotic plaques ^1, 8^. The removal of these senescent cells attenuated the onset and progression of cardiovascular diseases including atherosclerosis ^1, 8^.

The senescent cells acquire a pro-inflammatory phenotype, known as senescence- associated secretory phenotype (SASP) ^2^. These cells increase the expression and release of inflammatory cytokines, chemokines, cell adhesion molecules, and matrix metalloproteinase ^2, 4, 5, 9^. The SASP factors released from senescent cells affect the function of the neighboring cells and induce senescence in them ^10^. This paracrine process is called the bystander effect and exacerbates inflammation in the tissue surrounding senescent cells^10^. Moreover, SASP chemokines and adhesion molecules promote the recruitment of inflammatory cells into the vascular tissue ^10^. The SASP factors activate smooth muscle cells and infiltrated inflammatory cells such as monocytes and neutrophils, which then release pro- inflammatory molecules and MMPs, promoting inflammation and tissue remodeling in a feed- forward response ^11^, thus consequently resulting in the formation and progression of atherosclerotic plaques ^10^. In addition to tissue re-modulation, MMPs contribute to inflammation through their proteinase activity on inflammatory molecules ^11^. The lack of these SASP factors or their inhibition by blocking their receptors has been shown to protect against vascular diseases such as atherosclerosis and cerebral aneurysms (CAs) formation and rupture in experimental animal studies ^12–18^.

Oxidative stress induces endothelial cell dysfunction and senescence. Cardiovascular risk factors like ethanol, smoking, and hypertension produce oxidative stress, which subsequently induces DNA damage resulting in cellular senescence. Oxidative stress and oxidative stress- induced DNA damage can activate NF-κB, MAPKs, and mTOR pathways ^19^. These pathways are known to contribute to inflammation and cellular senescence ^9, 19–21^. Moreover, P38 regulates SASP factors at the mRNA level through NF-κB transcriptional activity ^9, 19^. Previous studies have suggested that blocking the activation of these pathways can dampen inflammation, senescence, and SASP ^20, 21^.

Colchicine is an anti-inflammatory drug that has been used for various diseases. Clinical and experimental animal studies have reported its protective effects against cardiovascular diseases. In animal experimental studies, colchicine mitigated atherosclerosis ^22, 23^, thrombosis ^24^, cardiac damage ^25^ and heart failure ^26^. In clinical trials, a daily dose of 0.5 mg of colchicine lowered the risks of cardiovascular events in patients ^27, 28^.

In the current study, we investigated the mechanisms, through which colchicine provides benefits against cardiovascular diseases. Here, we show that colchicine blocked oxidative stress-induced endothelial cell senescence and reversed a SASP via inhibiting NF-κB and MAPKs.

## Methods

### Cell culture

HUVECs were commercially obtained from Promocell (Heidelberg, Germany) and maintained in the endothelial cell medium (C-22010, Promocell, Heidelberg, Germany) supplemented with endothelial cell growth factors (C-39215, Promocell, Heidelberg, Germany) at 37 °C in a 95 % humidified atmosphere containing 5 % CO_2_. Cells were seeded in the T75 adherent cell culture flask after thawing. When the cells came to 90 % confluence, they were incubated with trypsin for 5 minutes at 37 °C, and then cells were used for experiments at passage 7. After 24 hours, the medium was changed and the new medium containing either H_2_O_2_ (300 µM), colchicine (50 nM), or H_2_O_2_ (300µM) combined with colchicine (50 nM) was added to the culture. Colchicine was purchased from Sigma-Aldrich (C3915).

### ROS Assay

ROS assay was performed using a Total Reactive Oxygen Species Assay Kit 520nM (Cat. Nr. 88-5930-74, Thermo Fisher Scientific) following the manufacturer’s instructions. Briefly, HUVECs were seeded in 96 well adhesion plates with the density of 2000 cells / cm2, then put into a 5% CO_2_, 95% humidity incubator for 24h. The next day, the medium was changed with a new medium containing 1x ROS Assay Stain stock solution and incubated for 1h. After that, either 300μM H_2_O_2_, 50 nM colchicine, or 300μM H_2_O_2_ combined with 50 nM colchicine were added to the medium already containing 1x ROS Assay Stain stock solution. The endothelial cell medium alone was added to the control cells. The measurements for ROS assay were performed with the Paradigm micro-plate reader after 2 h, 4 h, 6 h, 24 h 48 h of treatments.

### Immunofluorescence Staining

HUVECs were seeded in a 96-well plate (5000 cells/cm2). The medium was changed the next day with a new medium alone (for control) or a medium containing 300 µM H_2_O_2_, 50 nM colchicine, and a combination of 300 µM H_2_O_2_ and 50 nM colchicine. After two hours of treatment, immunofluorescence staining was performed. The cells were washed 3 times with PBS and fixed with 4% paraformaldehyde for 10 min, permeabilized with 0.2% Triton™ X- 100 for 10 min, and blocked with 5% bovine serum albumin (BSA) for 1 h at RT. The cells were incubated overnight at 4° C with primary antibody 8-OHDG (1:500, Cat. No. BSS-BS- 1278R, BIOSS, USA). The next day, the cells were washed three times with PBS and then labeled with a secondary antibody (1:1000, Alexa Fluor 488, ab150077, Abcam) for 1 h at RT. Nuclei were stained with DAPI (1ug/ml, 62248, Thermo Fisher Scientific) for 10 min. The images were captured at 20x magnification under a Leica DMi8 Inverted Microscope and the compatible LAS-X Life Science Microscope Software Platform.

### β-Galactosidase (β-Gal) staining

β-Gal staining was performed with a β-Galactosidase Reporter Gene Staining Kit (Sigma-Aldrich, St. Louis, USA), following manufacturer instructions. The endothelial cells were treated with endothelial cell medium alone (control group) or endothelial cell medium supplemented with either 300 µM H_2_O_2_ or 50 nm colchicine or 300 µM H_2_O_2_ combined with 50 nM colchicine. After 24h the cells were fixed in the fixation buffer provided with the β-Galactosidase Reporter Gene Staining Kit. The fixed samples were stained at 37 °C for 7 hours. After aspiration of the staining solution, the cells were overlaid with a 70 % glycerol solution and stored at 4 °C. The images were taken with a Leica DMi8 Inverted Microscope and the compatible LAS-X Life Science Microscope Software Platform. Image J was used for the counting of stained cells. β-Gal staining was performed with three biological triplicates.

### Western blot

For protein analysis, HUVECs at P7 were treated with endothelial cell medium alone (Control group) or endothelial cell medium supplemented with either 300 µM H_2_O_2_, 50 nM colchicine, or 300 µM H_2_O_2_ combined with 50 nM colchicine in biological triplicates. After 24 h of treatment, the total protein was extracted using RIPA buffer, and protein concentration was determined using the DC Protein Assay Kit (500-0116, Bio-Rad, Hercules, CA, USA) following the manufacturer’s instructions, and measured with the Paradigm micro-plate reader. Total protein (25 µg, in reducing conditions) was loaded on 12 % sodium dodecyl sulfate-polyacrylamide gel and run at 60 volts for 20 min, then continued with 110 Volts for 60 min, which was further transferred onto a polyvinylidene difluoride membrane at 250 mA for 120 min. The non-specific binding was blocked with 5 % BSA dissolved in 0.05 % TBST for 1 h. The membranes were incubated with primary antibodies (as reported in Supplementary Table 1) overnight at 4 °C on a shaking platform. The membranes were washed three times for 10 min with TBST and then incubated with secondary antibodies (as reported in Supplementary Table 1) for 1 h at room temperature. All primary antibodies were diluted in the blocking solution containing 5 % BSA. The secondary antibodies were diluted in TBST. The densitometry of immunoblotted bands was calculated with Image J (Version 1.53t, National Institutes of Health, Bethesda, MD, USA). All experiments were performed in triplicates.

### Quantitative PCR

For quantitative PCR (qPCR) analysis, HUVECs at P7 were treated with either endothelial cell medium alone (control group) or endothelial cell medium supplemented with either 300 µM H_2_O_2_ or 50 nM colchicine or 300 µM H_2_O_2_ combined with 50 nM colchicine in biological triplicates for 24 h. Total RNA was extracted using NucleoSpin RNA, Mini kit (740955.50, MACHEREY-NAGEL) following the manufacturer’s instructions. RNA (1.2 µg) was utilized to reverse transcript with M-MLV Reverse Transcriptase kit (M1701, Promega), Random Hexamer Primers (48190011, Thermo Fisher Scientific), and RiboLock RNase Inhibitor (EO0384, Thermo Fisher Scientific). qPCR was performed with AceQ SYBR qPCR Master Mix (Q111-03, Vayzme, Nanjing, China) on Bio-Rad CFX Connect Real-Time PCR System with an initial denaturation of 95 °C for 8 minutes, followed by 40 cycles of 95 °C for 15s, 58.9 °C for 30s, and 72 °C for 30s, followed by a melting curve. After normalizing to β-actin expression the relative mRNA expressions were calculated using the comparative ΔCT method. The primer sequences used in experiments are reported in Supplementary Table 2.

### Statistical Analysis

The data was analyzed by applying One-way ANOVA followed by Tukey’s test using Prism 9.5.0 (GraphPad Software Inc., San Diego, CA, USA). *P < 0.05 was considered significant.

## Results

### Colchicine inhibited H_2_O_2_ induced ROS generation and oxidative stress-induced DNA damage

To investigate the effect of colchicine on ROS generation, we treated endothelial cells with either 300 µM H_2_O_2_, 50 nM colchicine, or 300 µM H_2_O_2_ combined with 50 nM colchicine. The endothelial cell medium alone was used for untreated control. Colchicine inhibited ROS generation in H2O2-treated endothelial cells at all investigated time points (after 2 h: Control = 4.23 ± 0.21 arbitrary unit (a.u.), H_2_O_2_ = 99.83 ± 8.31 a.u., Colchicine = 4.41 ± 0.51 a.u., H_2_O_2_ + Colchicine = 4.31 ± 0.13 a.u., after 4 h: Control = 5.02 ± 0.22 a.u., H_2_O_2_ = 102.19 ± 6.94 a.u., Colchicine = 5.31 ± 0.45 a.u., H_2_O_2_ + Colchicine = 5.32 ± 0.16 a.u., after 6 h: Control = 6.60 ± 0.33 a.u., H_2_O_2_ = 104.66 ± 5.90 a.u., Colchicine = 6.91 ± 0.46 a.u., H_2_O_2_ + Colchicine = 6.79 ± 0.29 a.u., after 24 h: Control = 30.78 ± 2.27 a.u., H_2_O_2_ = 126.47 ± 9.28 a.u., Colchicine = 32.20 ± 1.26 a.u., H_2_O_2_ + Colchicine = 32.05 ± 0.304 a.u., after 48 h: Control = 46.46 ± 5.01 a.u., H_2_O_2_ = 135.57 ± 12.22 a.u., Colchicine = 47.55 ± 2.32 a.u., H_2_O_2_ + Colchicine = 49.97 ± 2.65 a.u., n = 3, ****p<0.0001, Figure 1A). Previously colchicine has been reported to inhibit ROS generation in platelets, endothelial cells, and macrophages^24, 29, 30^. ROS can cause DNA damage, growth arrest, and premature cellular senescence ^2^.

**Figure 1.**
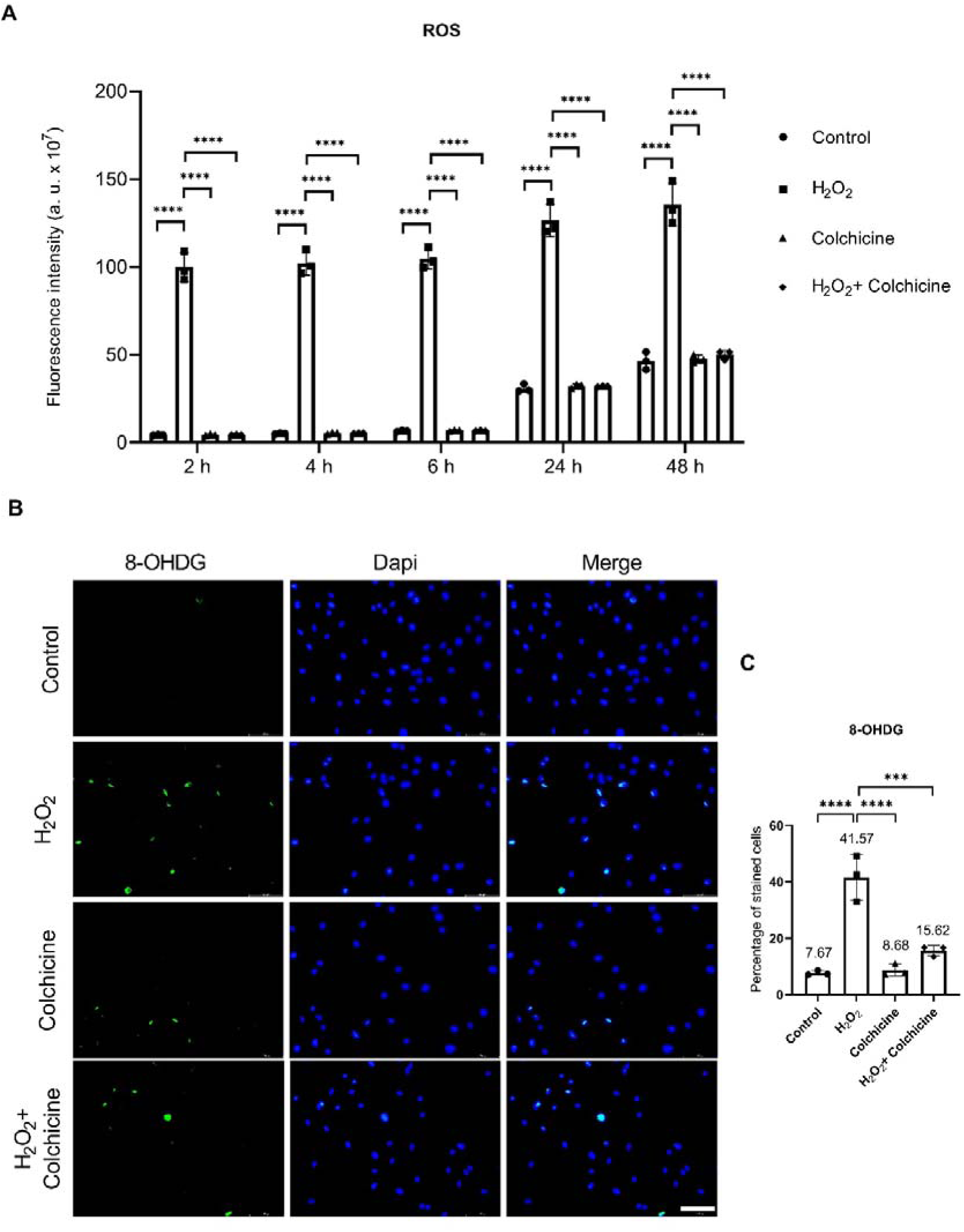
Colchicine abates ROS generation and oxidative-stress-induced DNA damage. (A) Colchicine inhibited ROS generation in endothelial cells treated with H_2_O_2_ at all investigated time points. (B) Immunofluorescence staining for oxidative stress-induced DNA damage marker 8-OHDG. (C) Colchicine averted oxidative-stress-induced DNA damage in endothelial cells exposed to H_2_O_2_-induced oxidative stress. The experiment was performed with biological triplicates. The data was analyzed by applying One-way ANOVA followed by Tukey’s test. Arbitrary unit (a.u.), error bars represent the standard deviation (SD), (**** p < 0.0001, *** p < 0.001), scale bar = 100 µm.

Next, we performed immunofluorescence staining for oxidative stress-induced DNA damage marker 8-OHDG in endothelial cells treated with similar conditions for 2 h. Colchicine ameliorated oxidative stress-induced DNA damage in endothelial cells, a percent of 8-OHDG positive cells (Control = 7.67 ± 0.8223 %, H_2_O_2_ = 41.57 ± 8.135 %, Colchicine = 8.68 ± 2.109 %, H_2_O_2_ + Colchicine = 15.62 ± 1.894 %, n = 3, ***p<0.001, ****p<0.0001, Figure 1B, C).

Previously, colchicine has been reported to reduce oxidative stress-induced DNA damage in ethanol-treated endothelial cells ^5^ and to provide anti-oxidative effects in endothelial cells and platelets via increasing the expression of anti-oxidative enzymes ^24, 30^. The oxidative stress and oxidative stress-induced DNA damage can activate NF-κB, MAPKs, and mTOR pathways, and can cause cellular senescence ^9, 19^.

### Colchicine inhibited oxidative-stress-induced senescence

To investigate whether colchicine can attenuate oxidative stress-induced senescence in endothelial cells, we subjected endothelial cells to 300 µM H_2_O_2_, colchicine 50 nM, or 300 µM H_2_O_2_ combined with 50 nM colchicine treatment for 24 h. The endothelial cell medium alone was used for untreated control. After 24 hrs, we performed β-gal staining. Colchicine obviated oxidative stress-induced senescence in endothelial cells, a percentage of β-gal positive cells (Control = 9.78 ± 0.74 %, H_2_O_2_ = 36.17 ± 5.74 %, Colchicine = 10.79 ± 1.31 %, H_2_O_2_ + Colchicine = 16.30 ± 3.17 %, n = 3, *** p < 0.001, **** p < 0.0001, Figure 2A, B). Oxidative stress, colchicine, or oxidative stress combined with colchicine did not alter the relative protein expression of DNA repair protein KU70 (Control = 1.00 ± 0.08, H_2_O_2_ = 0.90 ± 0.06, Colchicine = 1.03 ± 0.06, H_2_O_2_ + Colchicine = 0.94 ± 0.05, n = 3, Figure 2C, D). Both oxidative stress and colchicine reduced the relative protein expression of DNA repair protein KU80 (Control = 1.00 ± 0.08, H_2_O_2_ = 0.52 ± 0.04, Colchicine = 0.79 ± 0.08, H_2_O_2_ + Colchicine = 0.94 ± 0.02, n = 3, *p<0.05, **p<0.01, ***p<0.001, ****p<0.0001, Figure 2C, E) and Lamin B1 (Control = 1.00 ± 0.03, H_2_O_2_ = 0.64 ± 0.03, Colchicine = 0.87 ± 0.04, H_2_O_2_ + Colchicine = 0.88 ± 0.05, n = 3, *p<0.05, ***p<0.001, ****p<0.0001, Figure 2C, F). However, the relative protein expression of KU80 and Lamin B1 was significantly higher in endothelial cells treated with colchicine or colchicine combined with H_2_O_2_ than in the endothelial cells treated with H_2_O_2_ alone for 24 h. KU80 forms a heterodimer with KU70 and repairs double-strand DNA breaks through non-homologous end joining ^31^. Reduced levels of KU80 have been observed in senescent cells ^32^ and impaired KU80 protein expression has been shown to cause telomere shortening ^33^, which can consequently result in cellular senescence. The aging marker Lamin B1 maintains nuclear stability and reduced Lamin B1 protein levels resulted in misregulated non-homologous end joining and homologous repair of DNA leading to persistent DNA damage ^34^. Lamin B1 protein level is decreased in senescent cells ^35, 36^ due to its reduced stability ^35^. Loss of Lamin B1 protein expression can cause premature senescence ^36^. Moreover, colchicine ameliorated the relative protein expression of senescent marker P21 (Control = 1.001 ± 0.036, H_2_O_2_ = 2.097 ± 0.094, Colchicine = 1.060 ± 0.113, H_2_O_2_ + Colchicine = 1.398 ± 0.087, n = 3, **p<0.01, ****p<0.0001, Figure 2C, G). The quantitative analysis of P21 mRNA expression showed that P21 expression is regulated at the mRNA level (Control = 1.002 ± 0.0652, H_2_O_2_ = 3.973 ± 0.4731, Colchicine = 1.930 ± 0.2538, H_2_O_2_ + Colchicine = 3.083 ± 0.3428, n = 3, *p<0.05, ***p<0.001, ****p<0.0001, Figure 2H). P21 is an inhibitor of the cyclin-dependent kinase (CDK) and establishes indefinite growth arrest of senescent cells ^2^. The induction of P21 protein expression led to senescence in human HT1080 fibrosarcoma cells ^37^.

**Figure 2.**
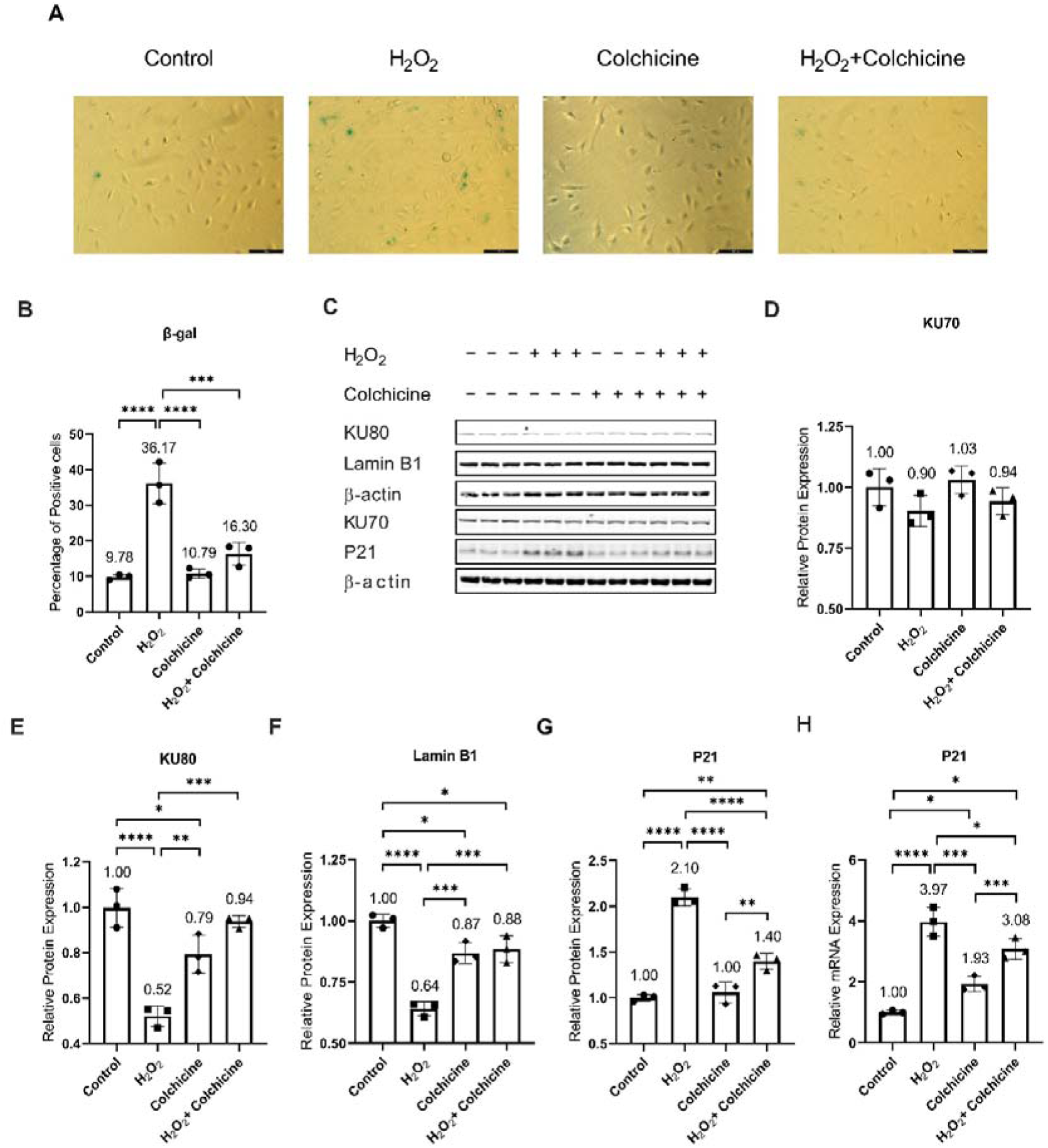
Colchicine subdues oxidative stress-induced senescence and restored the expression of senescence markers in endothelial cells. (A) β-gal staining of endothelial cells treated with 300 µM H_2_O_2_ or 50 nM colchicine, or a combination of 300 µM H_2_O_2_ and 50 nM colchicine for 24 h. Endothelial medium alone was used for untreated control cells. (B) Colchicine averted senescence in endothelial cells exposed to oxidative stress. (D) H_2_O_2_, colchicine, or H_2_O_2_ combined with colchicine did not affect the relative protein expression of DNA repair protein KU70. (E) Colchicine improved the expression of DNA repair protein KU80. (F) Colchicine rescued the relative protein expression of the aging-associated marker Lamin B1 and (G) attenuated the relative protein expression of the senescence marker P21. The experiment was performed with biological triplicates. The data was analyzed by applying One-way ANOVA followed by Tukey’s test. Error bars represent SD, (****<0.0001, *** p < 0.001, ** p < 0.01, * p < 0.05), scale bar = 100 µm.

These findings suggest that colchicine inhibited oxidative stress-induced senescence (Figure 2A, B) via improving KU80 (Figure 2E) and Lamin B1 (Figure 2F) protein expression and attenuating P21 (Figure 2G, H) expression at mRNA and protein levels. The expression of senescence markers and DNA repair proteins is regulated by NF-κB, MAPKs, and mTOR pathways and these pathways are known to contribute to cellular senescence ^2, 9, 19^. Therefore, we performed a Western blot for protein analysis to investigate if colchicine regulates the activation of these pathways in oxidative stress-induced endothelial cells.

### Pathway analysis

The protein analysis showed that both oxidative stress and colchicine neither alone nor in combination changed the relative protein expression of NF-κB subunit P65 at both investigated time points after H_2_O_2_ treatment (Relative protein expression of P65 after 30 min: Control = 1.001 ± 0.0497, H_2_O_2_ = 0.9051 ± 0.03, Colchicine = 1.007 ± 0.029, H_2_O_2_ + Colchicine = 1.021 ± 0.055; after 2 h: Control = 1.000 ± 0.1116, H_2_O_2_ = 0.8558 ± 0.0153, Colchicine = 1.006 ± 0.0593, H_2_O_2_ + Colchicine = 1.034 ± 0.0206, n = 3, *p < 0.05, Figure 3A, B). Colchicine inhibited activation of NF-κB in endothelial cells exposed to H_2_O_2_ for 30 min (Relative protein expression of p-P65: Control = 0.9995 ± 0.017, H_2_O_2_ = 2.426 ± 0.4105, Colchicine = 0.8756 ± 0.016, H_2_O_2_ + Colchicine = 1.441 ± 0.22, n = 3, **p < 0.01, ***p < 0.001, Figure 3A, C) and at the same time point p-P65/P65 ratio (Control = 0.9999 ± 0.041, H_2_O_2_ = 2.673 ± 0.3642, Colchicine = 0.87 ± 0.023, H_2_O_2_ + Colchicine = 1.407 ± 0.146, n = 3, ***p < 0.001, ****p < 0.0001, Figure 3D) was significantly reduced by colchicine in H_2_O_2_- treated endothelial cells. The relative protein expression of NF-κB subunit p-P65 (Control = 1.000 ± 0.2173, H_2_O_2_ = 1.047 ± 0.0798, Colchicine = 1.038 ± 0.1269, H_2_O_2_ + Colchicine = 1.277 ± 0.0519, n = 3, *p < 0.05, Figure 3A, C) and p-P65/P65 ratio (Control = 0.9925 ± 0.1008, H_2_O_2_ = 1.224 ± 0.099, Colchicine = 1.029 ± 0.0652, H_2_O_2_ + Colchicine = 1.235 ± 0.0345, n = 3, *p < 0.05, Figure 3D) did not change in endothelial cells treated with H_2_O_2_, colchicine, or H_2_O_2_ and colchicine combined for 2 h.

**Figure 3.**
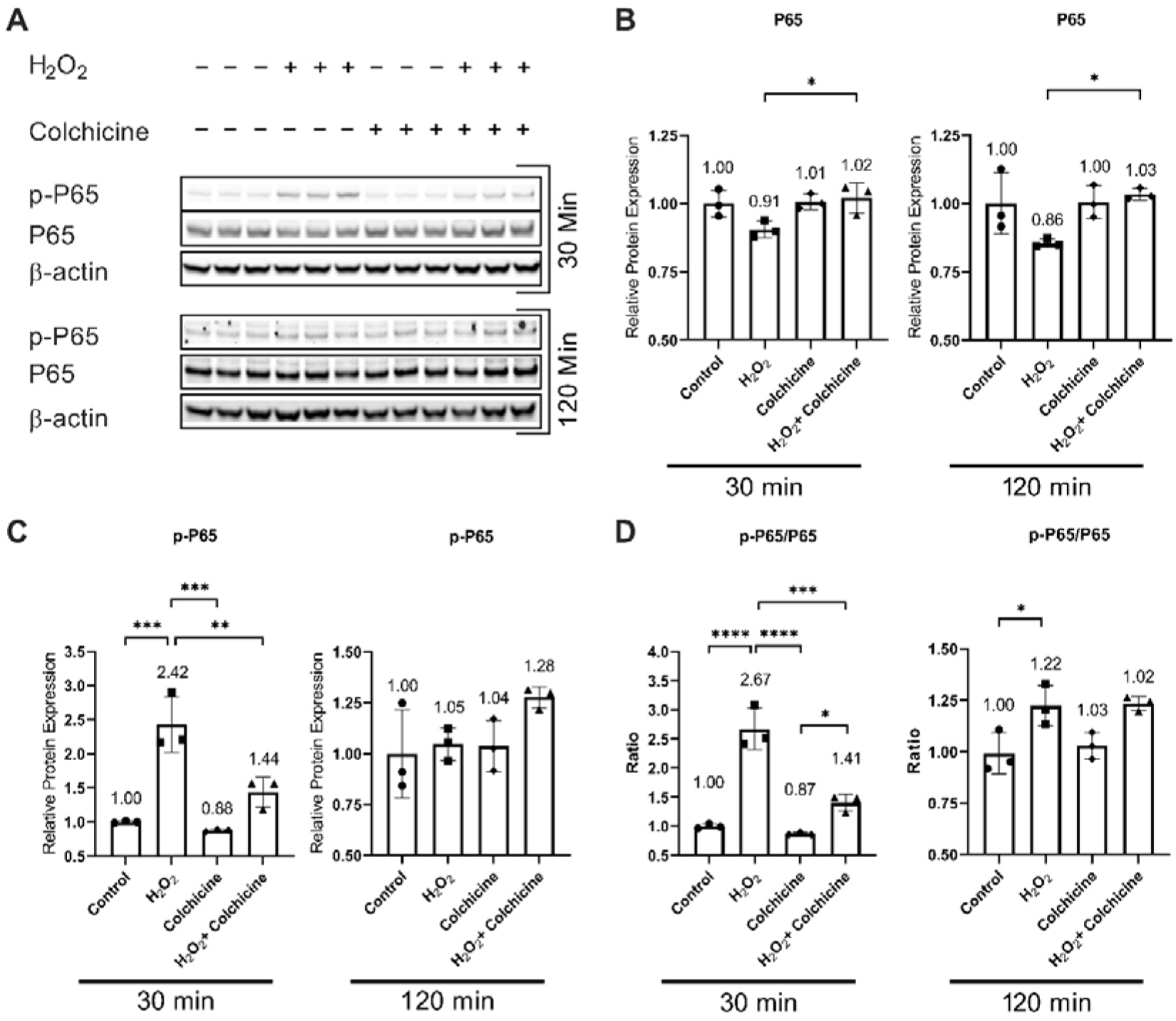
Colchicine inhibits NF-κB activation: (A) Protein expression of NF-κB subunits in endothelial cells exposed to 300 mM H_2_O_2_, 50 nM colchicine, or 300 mM H_2_O_2_ combined with 50 nM colchicine. The control was treated with endothelial cell medium alone. (B) Oxidative stress and colchicine alone or in combination did not affect the relative protein expression of NF-κB subunit P65 when compared to untreated control. (C) Oxidative stress increased the relative protein expression of NF-κB subunit p-P65 after 30 minutes of exposure, which was inhibited by colchicine. Both oxidative stress and colchicine neither alone nor in combination alter the relative protein expression of NF-κB subunit p-P65 after 120 min exposure. (D) Colchicine reduced p-P65/P65 ratio in endothelial cells exposed to H_2_O_2_ for 30 min. The H_2_O_2_ treatment for 120 min increased p-P65/P65 ratio in endothelial cells. The experiment was performed with biological triplicates. The data was analyzed by applying One-way ANOVA followed by Tukey’s test. Error bars represent the SD (*** p < 0.001, ** p < 0.01, * p < 0.05).

NF-κB is known to promote inflammation and contribute to cellular senescence *in vitro* and *in vivo* ^9, 10, 38^. NF-κB activation impairs KU80 protein expression that subsequently results in telomere shortening ^39^, which can lead to cellular senescence. In response to DNA damage, NF-κB increased the expression of P21 ^40^ leading to cell cycle arrest and senescence ^2, 38^. This suggests that colchicine via blocking NF-κB activation can suppress P21 expression (Figure 1G, H) and improve KU80 expression (Figure 1E), resulting in reduced endothelial cell senescence (Figure 1A, B). Moreover, NF-κB regulates SASP responses in dysfunctional and senescent endothelial cells ^9, 10^, where the transcriptional activity of NF-κB in senescent cells is regulated by MAPKs ^19^.

Oxidative-stress-induced DNA damage can activate MAPKs ^19^. Protein analysis showed that H_2_O_2_ induced oxidative-stress-activated MAPKs (Figure 4). Colchicine inhibited oxidative-stress-induced activation of P38 (relative protein expression of p-P38 after 30 min: Control = 0.9975 ± 0.095, H_2_O_2_ = 4.423 ± 0.872, Colchicine = 0.805 ± 0.173, H_2_O_2_ + Colchicine = 1.652 ± 0.121; after 2 h: Control = 1.000 ± 0.0669, H_2_O_2_ = 1.857 ± 0.2346, Colchicine = 1.213 ± 0.1798, H_2_O_2_ + Colchicine = 1.217 ± 0.096, n = 3, **p < 0.01, ***p < 0.001, ****p < 0.0001, Figure 4A, B) and ERK (Relative protein expression of p-ERK after 30 min: Control = 1.001 ± 0.1285, H_2_O_2_ = 3.275 ± 0.3148, Colchicine = 0.668 ± 0.1129, H_2_O_2_ + Colchicine = 1.048 ± 0.1542; after 2 h: Control = 0.9999 ± 0.1663, H_2_O_2_ = 3.275 ± 0.1169, Colchicine = 1.021 ± 0.2034, H_2_O_2_+Colchicine = 0.9414 ± 0.1851, n = 3, ****p < 0.0001, Figure 4A, C) in endothelial cells at both investigated time points after H_2_O_2_ treatment. Both oxidative stress and colchicine neither alone nor in combination had an effect on the activation of JNK (Relative protein expression of p-JNK after 30 min: Control = 0.9971 ± 0.017, H_2_O_2_ = 1.012 ± 0.027, Colchicine = 1.012 ± 0.02, H_2_O_2_ + Colchicine = 1.030 ± 0.016; after 2 h: Control = 1.000 ± 0.028, H_2_O_2_ = 0.938 ± 0.019, Colchicine = 0.9002 ± 0.0378, H_2_O_2_ + Colchicine = 0.9098 ± 0.0215, n = 3, *p < 0.05, Figure 4A, D) at both investigated time points after H_2_O_2_ exposure.

**Figure 4.**
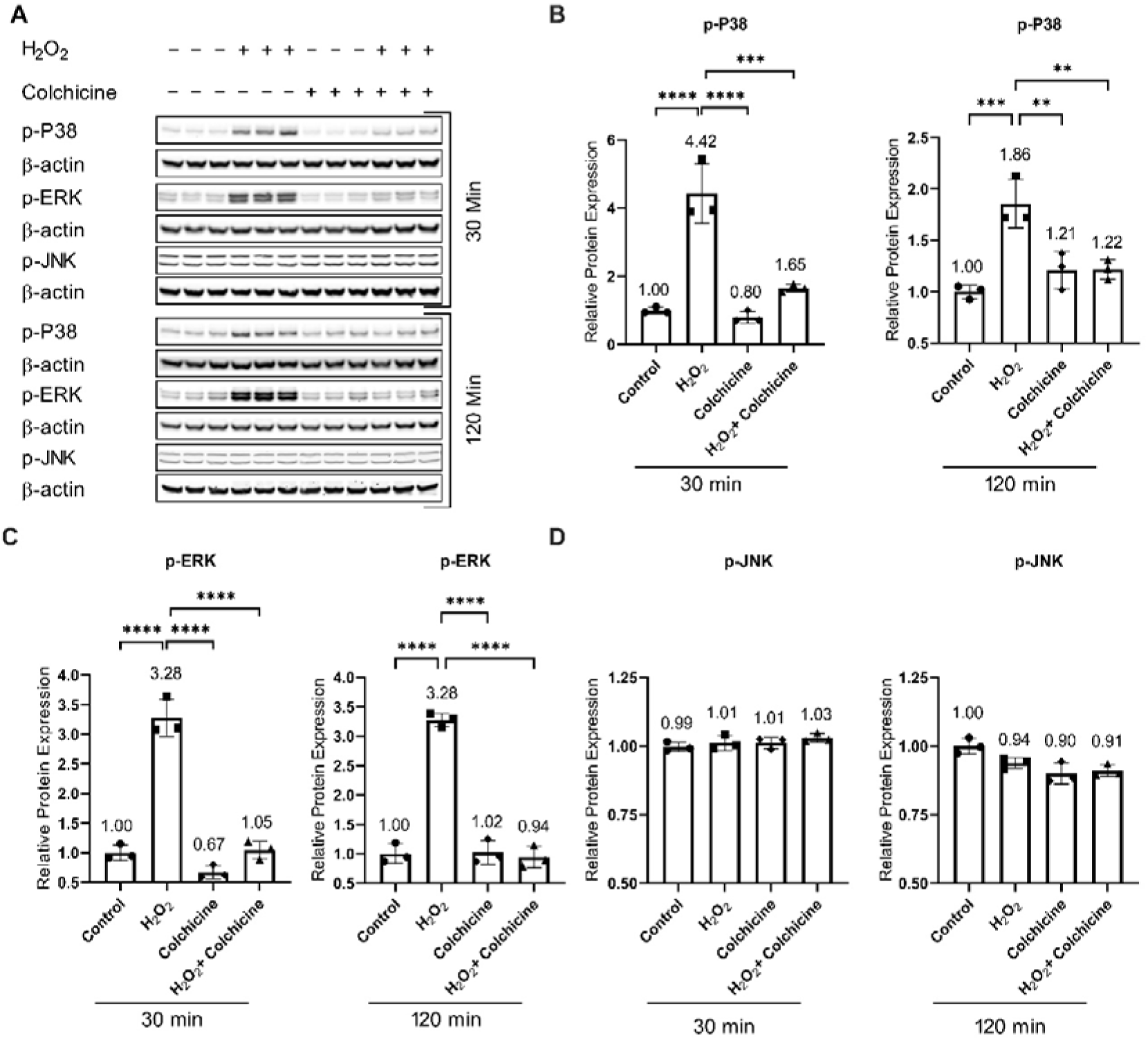
Colchicine inhibits oxidative stress-induced P38 and ERK activation: (A) Protein expression of MAPKs in endothelial cells exposed to oxidative stress, colchicine, or oxidative stress combined with colchicine at different time points. After 30 min and 120 min treatment of H_2_O_2_, colchicine inhibited the activation of (B) P38 and (C) ERK. (D) Both H_2_O_2_ and colchicine neither alone nor in combination alter the relative protein expression of p-JNK. β- actin was used as a loading control. The experiment was performed with biological triplicates. The data was analyzed by applying One-way ANOVA followed by Tukey’s test. Error bars represent the SD, ****p < 0.0001, *** p < 0.001, ** p < 0.01, and * p < 0.05.

MAPKs regulate the protein expression of cell cycle arrest proteins and SASP factors ^19^. P38 increases the expression, stabilization, and promoter activity of P53, resulting in increased P21 protein expression ^41, 42^. P38 increases the cytoplasmic accumulation of HuR via phosphorylating it ^43^. HuR binds to P21 mRNA and increases its stability, consequently increasing P21 protein expression ^43^. Blocking senescent signals via inhibiting P38 can mitigate Lamin B1 loss ^35^. ERK also promotes the transcription of P21 ^19^. Previous studies have shown that inhibiting P38 activation can subdue cellular senescence ^44^. Taken together, it seems plausible that colchicine (Figure 2A, B) via inhibiting P38 (Figure 4B) can improve Lamin B1 protein expression (Figure 2F) and by blocking P38 (Figure 4B) and ERK (Figure 4C) pathways can subdue P21 mRNA and protein expression (Figure 1G, H), and thus suppresses endothelial cell senescence (Figure 2A, B).

Protein analysis showed that colchicine inhibited oxidative stress-induced activation of AKT (Relative protein expression of p-AKT after 30 min: Control = 0.100 ± 0.009, H_2_O_2_ = 0.2547 ± 0.036, Colchicine = 0.079 ± 0.002, H_2_O_2_ + Colchicine = 0.1377 ± 0.01; after 2 h: Control = 1.000 ± 0.0559, H_2_O_2_ = 1.846 ± 0.0595, Colchicine = 1.040 ± 0.1119, H_2_O_2_ + Colchicine = 1.034 ± 0.1325, n = 3, ***p < 0.001, ****p < 0.0001, Figure 5A, B) at both investigated time points. Oxidative stress and colchicine neither alone nor in combination affected the relative protein expression of p-mTOR (Control = 1.003 ± 0.097, H_2_O_2_ = 1.300 ± 0.046, Colchicine = 1.216 ± 0.144, H_2_O_2_ + Colchicine = 1.155 ± 0.191, n = 3, *p < 0.05, Figure 5A, C) in endothelial cells treated for 30 min. Oxidative stress increased the relative protein expression of p-mTOR (after 2 h: Control = 1.000 ± 0.132, H_2_O_2_ = 1.444 ± 0.095, Colchicine = 1.370 ± 0.0881, H_2_O_2_ + Colchicine = 1.277 ± 0.1230; after 24 h: Control = 0.993 ± 0.178, H_2_O_2_ = 1.452 ± 0.069, Colchicine = 1.318 ± 0.013, H_2_O_2_ + Colchicine = 1.110 ± 0.137, n = 3, *p < 0.05, **p < 0.01, Figure 5A, C) in endothelial cells exposed to H_2_O_2_ for 2 and 24h. Colchicine inhibited the relative protein expression of p-mTOR in endothelial cells treated with oxidative stress for 24 h (Figure 5C). Both oxidative stress and Colchicine neither alone nor in combination alter the relative protein expression of p-S6 (after 30 min: Control = 1.000 ± 0.0140, H_2_O_2_ = 0.9447 ± 0.017, Colchicine = 1.02 ± 0.0187, H_2_O_2_ + Colchicine = 1.027 ± 0.032; after 24 h: Control = 0.998 ± 0.024, H_2_O_2_ = 0.941 ± 0.064, Colchicine = 1.036 ± 0.037, H_2_O_2_ + Colchicine = 1.053 ± 0.084, n = 3, *p < 0.05, **p < 0.01, Figure 5A, D) after 30 min and 24 h treatments when compared to untreated control. Colchicine did not affect oxidative stress-induced relative protein expression of p-S6 (Control = 1.000 ± 0.0769, H_2_O_2_ = 1.402 ± 0.0379, Colchicine = 1.319 ± 0.0661, H_2_O_2_ + Colchicine = 1.259 ± 0.0681, n = 3, **p < 0.01, ***p < 0.001, Figure 5A, D) in endothelial cells treated with H_2_O_2_ for 2 h. The 30 min combined treatment of oxidative stress and colchicine increased the relative protein expression of p-4-EBP1 (Control = 1.000 ± 0.173, H_2_O_2_ = 0.964 ± 0.112, Colchicine = 1.264 ± 0.076, H_2_O_2_ + Colchicine = 1.57 ± 0.159, n = 3, **p < 0.01, Figure 5A, E) than untreated control and oxidative treated stress alone. Rapamycin inhibition of the mTOR pathway in most cell types inhibits mTOR and its downstream signaling molecule S6 without affecting mTOR downstream signaling molecule 4-EBP-1 ^45^. Here, unlike rapamycin inhibition of mTOR pathway, colchicine recovered the relative protein expression of p-4EBP-1 in endothelial cells treated with oxidative stress for 2 h (Control = 1.000 ± 0.012, H_2_O_2_ = 0.774 ± 0.0466, Colchicine = 0.9912 ± 0.0729, H_2_O_2_ + Colchicine = 0.9724 ± 0.0291, n = 3, **p < 0.01, Figure 5A, E) and showed a trend of its increase after 24 h (Control = 1.000 ± 0.188, H_2_O_2_ = 0.869 ± 0.011, Colchicine = 0.802 ± 0.065, H_2_O_2_ + Colchicine = 1.101 ± 0.124, n = 3, *p < 0.05, Figure 5A, E).

**Figure 5.**
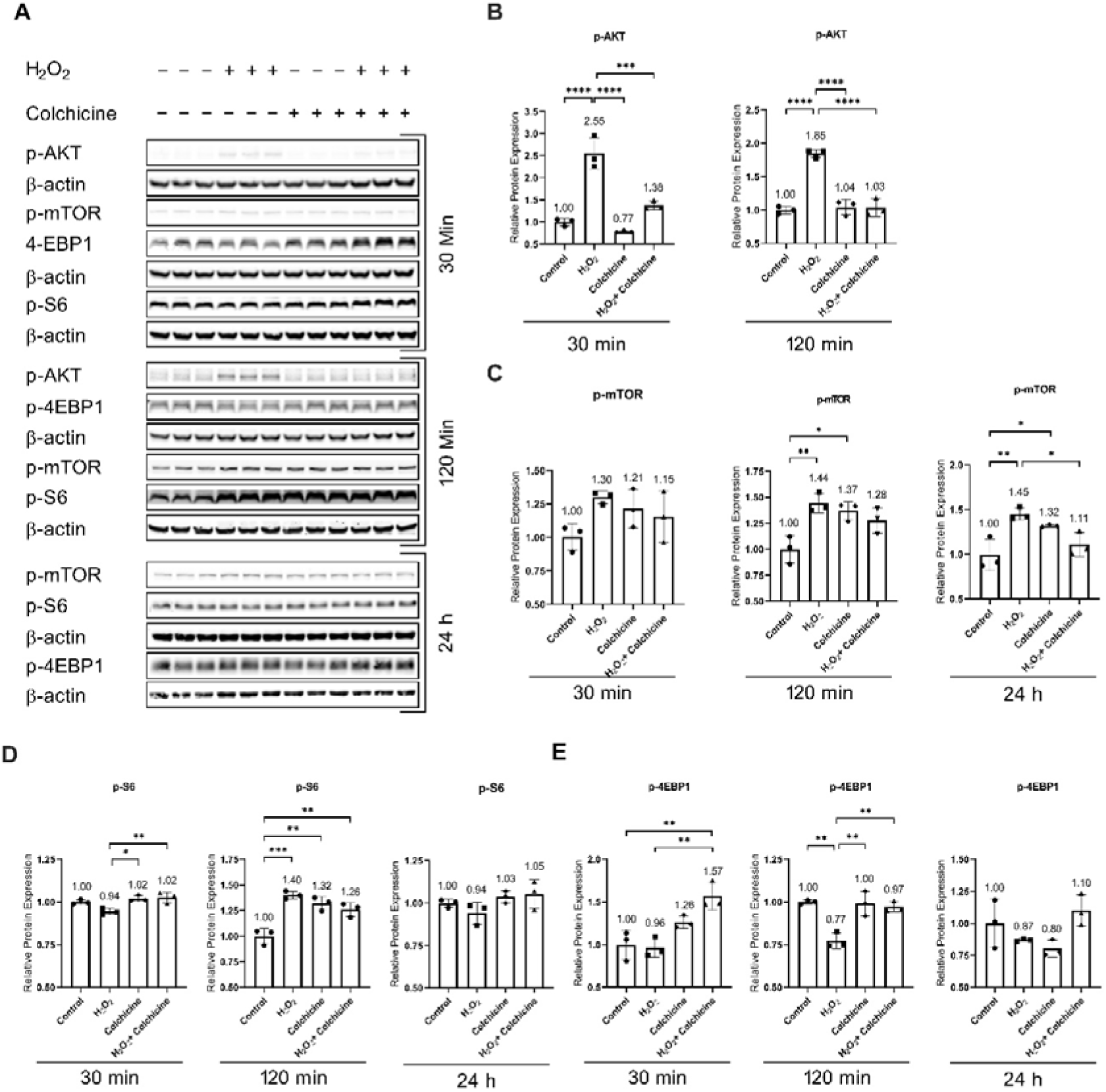
Colchicine modulates mTOR pathway activation: (A). Western blot showing protein expression of p-AKT, p-mTOR, p-4EBP1, and P-S6. (B) Colchicine subdued AKT activation in endothelial cells exposed to oxidative stress for 30 min and 120 min. (C) Neither of the treatment affected the relative protein expression of p-mTOR after 30 min. Both colchicine and oxidative stress increased the relative protein expression of p-mTOR after 120 min and 24 h. The relative protein expression of p-mTOR was lower in endothelial cells treated with colchicine combined with oxidative stress than in endothelial cells treated with oxidative stress alone for 24 h. (D) Neither oxidative stress nor colchicine nor oxidative stress combined with colchicine affected the relative protein expression of p-S6 in endothelial cells after 30 min and 24 h treatment. Oxidative stress, colchicine, and oxidative stress combine with colchicine increased the relative expression of p-S6 after 120 min treatment. (E) Colchicine combined with oxidative stress increased the relative protein expression of p- 4EBP1 as compared to control after 30 min treatment. Colchicine recovered the relative protein expression of 4-EBP1 in endothelial cells after 120 min treatment with H_2_O_2_. Neither of the treatment affected the relative protein expression of 4-EBP-1 after 24 h treatment. β- actin was used as a loading control. The experiment was performed with biological triplicates. The data was analyzed by applying One-way ANOVA followed by Tukey’s test. Error bars represent the SD, (*** p < 0.001, ** p < 0.01, and * p < 0.05).

mTOR is known to play a role in aging and senescence and its inhibition increases longevity and delays senescence ^20^. mTOR negatively regulates autophagy, resulting in accumulated damaged proteins and organelles, which can lead to the progression of cellular senescence^21, 46^. Blocking mTOR activation can improve mitochondrial function and reduce ROS levels ^21^, which can consequently provide protection against cellular senescence.

These findings indicate that colchicine inhibited oxidative stress-induced senescence (Figure 2) via inhibiting the activation of NF-κB, MAPKs, and mTOR pathway (Figures 3, 4, and 5). Because the activation of these pathways in senescent cells regulates the expression of SASP factors ^2, 9, 10, 19^, therefore, we investigated the effect of colchicine on the regulation of mRNA expression of SASP factors in endothelial cells treated with oxidative stress.

### Colchicine ameliorated oxidative stress-induced SASP in endothelial cells

Senescent cells increase the expression of inflammatory cytokines, chemokines, and cell adhesion molecules ^2, 4, 5^. Our results showed that oxidative stress increased the relative mRNA expression of IL-1β, IL-6, MCP-1, ICAM-1, and E-Selectin. Colchicine averted oxidative-stress-induced relative mRNA expression of cytokines: IL-1β (Control = 1.099 ± 0.5574, H_2_O_2_ = 2.026 ± 0.097, Colchicine = 0.4057 ± 0.1344, H_2_O_2_ + Colchicine = 1.149 ± 0.3112, n = 3, *p < 0.05, **p < 0.01, Figure 6A), IL-6 (Control = 1.005 ± 0.1179, H_2_O_2_ = 2.308 ± 0.2776, Colchicine = 1.741 ± 0.2189, H_2_O_2_ + Colchicine = 1.672 ± 0.0984, n = 3, *p <0.05, **p < 0.01, ***p < 0.001, Figure 6B), chemokine: MCP-1 (Control = 1.051 ± 0.3659, H_2_O_2_ = 1.871± 0.1475, Colchicine = 1.129 ± 0.1863, H_2_O_2_ + Colchicine = 0.3759 ± 0.1323, n = 3, *p <0.05, **p < 0.01, ***p < 0.001, Figure 6C), cell adhesion molecules: ICAM-1 (Control = 1.023 ± 0.2560, H_2_O_2_ = 2.260 ± 0.3143, Colchicine = 1.549 ±0.0639, H_2_O_2_ + Colchicine = 1.536 ± 0.1936, n = 3, *p <0.05, ***p < 0.001, Figure 6E) and E- selectin (Control = 1.029 ± 0.3132, H_2_O_2_ = 2.073 ± 0.0918, Colchicine = 0.5603 ± 0.0666, H_2_O_2_ + Colchicine = 0.6754 ± 0.2685, n = 3, **p < 0.01, ***p < 0.001, Figure 6G). The relative mRNA expression of IL-8 was significantly lower in endothelial cells treated with oxidative stress combined with colchicine than in endothelial cells treated with oxidative stress or colchicine alone (Control = 1.017 ± 0.2194, H_2_O_2_ = 1.367 ± 0.2210, Colchicine = 1.464 ± 0.180, H_2_O_2_ + Colchicine = 0.7314 ± 0.0211, n = 3, *p <0.05, Figure 6D). The mRNA expression of VCAM-1 was significantly higher in endothelial cells treated with oxidative stress combined with colchicine than untreated controls VCAM-1 (Control = 1.010 ± 0.1789, H_2_O_2_ = 1.301 ± 0.1466, Colchicine = 0.7173 ± 0.0827, H_2_O_2_ + Colchicine = 1.419 ± 0.1095, n = 3, *p <0.05, **p < 0.01, Figure 6F).

**Figure 6.**
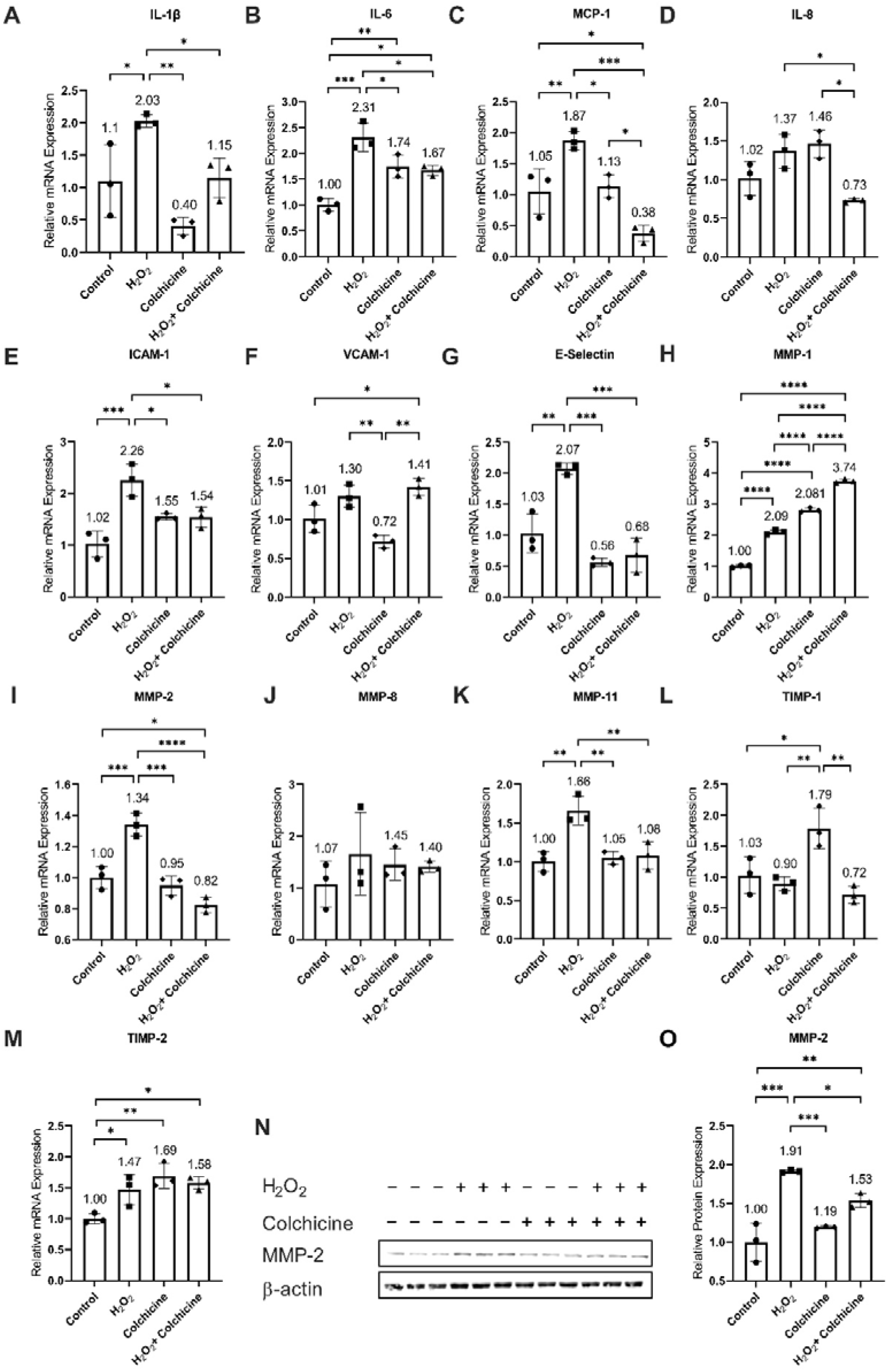
Colchicine mitigates the relative mRNA expression of SASP factors. Colchicine abated the relative mRNA expression of (A) IL-1β, (B) IL-6, (C) MCP-1, (E) ICAM-1, (G) E-selectin, (I) MMP2 and (K) MMP-11 in endothelial cells treated with H_2_O_2_ for 24 h. Both H_2_O_2_- induced oxidative stress and colchicine neither alone nor in combination altered the relative mRNA expression of (D) IL-8 and (J) MMP-8 when compared to untreated control. (F) Colchicine did not lower the oxidative stress-induced relative mRNA expression of VCAM-1 in endothelial cells. The 24 h treatment with H_2_O_2_, colchicine, or H_2_O_2_ combined with colchicine increased the relative mRNA expression of (H) MMP-1 and (M) TIMP-2. (L) Colchicine treatment for 24 h increased the relative mRNA expression of TIMP-1 in endothelial cells. (N) Western blot showing protein expression of MMP-2 in endothelial cells treated with H_2_O_2_ or colchicine or H_2_O_2_ combined with colchicine for 24 h. (O) Colchicine reduced the relative protein expression of MMP-2 in endothelial cells treated with H_2_O_2_. β- actin was used as a loading control. qPCR data are the mean of three independent technical replicates and WB data are the mean of the biological triplicates. The data was analyzed by applying One-way ANOVA followed by Tukey’s test. Error bars represent the SD, *** p < 0.001, ** p < 0.01, and * p < 0.05).

In addition to inflammatory markers, senescent cells increase the expression and release of MMPs ^4, 5, 10^, which consequently contributes to the initiation and progression of cardiovascular diseases ^10, 47^. We performed qPCR to investigate the effects of colchicine on the relative mRNA expression of MMPs in endothelial cells exposed to H_2_O_2_ for 24 h. H_2_O_2_, colchicine and H_2_O_2_ combined with colchicine increased the relative mRNA expression of MMP-1 (Control = 1.001 ± 0.0373, H_2_O_2_ = 2.096 ± 0.069, Colchicine = 2.810 ± 0.0643, H_2_O_2_ + Colchicine = 3.748 ± 0.060, n = 3, ***p < 0.001, Figure 6H) and TIMP-2 (Control = 1.002 ± 0.083, H_2_O_2_ = 1.471 ± 0.2438, Colchicine = 1.689 ± 0.2042, H_2_O_2_ + Colchicine = 1.578 ± 0.1015, n = 3, *p <0.05, **p < 0.01, Figure 6M). Colchicine reduced the relative mRNA expression of MMP-2 (Control = 1.002 ± 0.0728, H_2_O_2_ = 1.340 ± 0.0733, Colchicine = 0.950 ± 0.0627, H_2_O_2_ + Colchicine = 0.8241 ± 0.0484, n = 3, *p <0.05, ***p < 0.001, ****p < 0.0001, Figure 6I) and MMP-11 (Control = 1.005 ± 0.1287, H_2_O_2_ = 1.658 ± 0.1862, Colchicine = 1.051 ± 0.0813, H_2_O_2_ + Colchicine = 1.082 ± 0.1764, n = 3, **p < 0.01, Figure 6K). Both H_2_O_2_ and colchicine neither alone nor in combination affected the mRNA expression of MMP-8 (Control = 1.071 ± 0.4426, H_2_O_2_ = 1.656 ± 0.7994, Colchicine = 1.447 ± 0.3067, H_2_O_2_ + Colchicine = 1.409 ± 0.1112, n = 3, *p < 0.05, Figure 6J). Colchicine decreased the relative protein expression of MMP-2 (Control = 0.997 ± 0.25, H_2_O_2_ = 1.911±0.016, Colchicine = 1.192±0.014, H_2_O_2_+Colchicine = 1.534±0.089, n = 3, *p <0.05, **p < 0.01, ***p < 0.001, Figure 6N, O) in endothelial cells treated with H_2_O_2_.

The expression of these SASP factors is regulated by P38 through the transcriptional activity of NF-κB ^19^. Agreeing with previously reported findings, our study showed that colchicine inhibited the activation of NF-κB, P38, and ERK pathways ^5^. The data suggest that via inhibiting the activation of NF-κB (Figure 3), P38 (Figure 4), and ERK (Figure 4) pathways colchicine suppressed oxidative stress-induced mRNA expression of SASP factors (Figure 6) in endothelial cells.

## Discussion

Endothelial cell dysfunction and senescence are known to contribute to cardiovascular diseases ^1^. These cells increase the expression and release of pro-inflammatory molecules and matrix metalloproteinase ^4, 5^, which have a causal relationship with cardiovascular diseases including atherosclerosis, thrombosis, aneurysm pathophysiology, stroke, and heart infarct among others ^10, 47^. In the current study, we used colchicine to ameliorate oxidative stress-induced dysfunction and senescence in endothelial cells. Colchicine inhibited ROS generation, subdued DNA damage, ameliorated endothelial cell senescence, impeded the activation of NF-κB, MAPKs, and mTOR pathways, and attenuated the expression of SASP-associated pro-inflammatory molecules and MMPs in endothelial cells treated with oxidative stress.

Endothelial cell dysfunction and senescence have a potential link between inflammation and aging, both of which have been implicated in cardiovascular diseases ^1, 10^. The senescent cells accumulated in different organs and tissues exerted negative effects on health- and life- span and their abolition protected against age-related diseases and increased health- and life-span in animal studies ^6, 7, 48^. Previous studies have shown the accumulation of senescent cells in atherosclerotic lesions ^1, 8^ and colchicine by inhibiting cellular senescence (Figure 2) can prevent the progression of atherosclerosis and can promote atherosclerotic plaque stability ^1, 8, 10^.

Our pathway analysis showed that colchicine inhibited NF-κB, MAPKs, and mTOR activation (Figures 3, 4, 5), which agrees with the previously reported findings ^5^. The previously published reports implicated the activation of NF-κB, MAPKs, and mTOR pathways in the progression and rupture of atherosclerotic lesions and CAs ^49–53^. The activation of NF-κB and MAPKs has been detected in human and animal atherosclerotic lesions and CA walls ^49–51^. The inhibition of NF-κB activation in rats and NF-κB subunit P50 deficiency in mice prevented CAs formation ^49^ and blocking nuclear import of NF-κB in mice reduced atherosclerotic lesion size ^50^. Moreover, these approaches inhibited macrophage infiltration into the arterial walls ^49, 50^, increased anti-inflammatory (M2-type) macrophages in atherosclerotic lesions ^50^ and lowered *in vivo* and *in vitro* the mRNA expression of MCP-1, ICAM-1, VCAM-1, MMP-2, MMP-9, IL-1β, IL-6, TNF-α and iNOS ^49, 50^. The phosphorylation of MAPKs correlated with CAs size and was higher in ruptured CAs than un-ruptured CAs ^51^, which suggests MAPKs role in the progression and rupture of CAs. Macrophage autophagy was increased *in vitro* and *in vivo* by AKT inhibitor, mTOR inhibitor, or mTOR-siRNA and resulted in macrophage clearance *in vivo* ^52^. AKT inhibitor, mTOR inhibitor, or mTOR-siRNA decreased plaque vulnerability and reduced atherosclerotic plaque rupture ^52^. In addition, the animals treated orally with rapamycin an inhibitor of mTOR had a significantly thicker fibrous cap of the abdominal aortic plaque and a lower rupture rate ^53^. Rapamycin lowered serum levels and the percentage of cells stained positively for MCP-1, MMP-1, MMP-2, MMP-3, MMP-9, MMP- 12, and P-selectin in the abdominal plaques ^53^. In addition to that, the mRNA expression of MCP-1, MMP-1, MMP-2, MMP-3, MMP-9, MMP-12, and P-selectin in the abdominal plaques was significantly lower in the rapamycin-treated group ^53^. PRAS40 overexpression and treatment of Torin1 an inhibitor of mTOR reduced THP-1 cells adhesion to TNF-α treated endothelial cells and inhibition of mTOR activation by PRAS40 and Torin1 attenuated TNF-α induced upregulation of ICAM-1 and VCAM-1^54^. Taken together, it can be suggested that colchicine via blocking NF-κB (Figure 3), MAPKs (Figure 4), and mTOR (Figure 5) pathways can protect against cardiovascular diseases including atherosclerosis and CAs ^49–53^.

Senescent endothelial cells increase the expression of SASP factors (Figure 6) ^4, 5, 10^, contributing to cardiovascular diseases via multiple mechanisms ^10^. The enhanced expression and release of cytokines, chemokines, and cell adhesion proteins by these senescent endothelial cells causes tissue infiltration of monocytes, neutrophils, and platelets. The infiltration, accumulation, and activation of neutrophils, monocytes, and platelets contribute heavily to atherosclerosis and thrombosis ^10^. Colchicine reduced the activation of monocytes ^23^ and hindered the adhesion of monocytes to endothelial cells by suppressing the expression of VCAM-1 and ICAM-1 ^24^. In experimental animal studies, colchicine dampened the infiltration and recruitment of neutrophils and monocytes into the atherosclerotic plaques ^22, 23^ and infarct area of myocardium after myocardial ischemia ^25^. Colchicine *in vitro* and *in vivo* ameliorated platelet activation and inhibited platelet-platelet, platelet-monocyte, and platelet-neutrophil aggregation ^24, 29, 55^ and thus can protect against atherosclerosis and thrombosis ^24^. Moreover, colchicine *in vitro* mitigated the mRNA and protein expression of TNF-α, IL-1β, IL-6, IL-18, MCP-1, ICAM-1 and VCAM-1 (Figure 6) ^5, 24, 30^ and *in vivo* ameliorated their mRNA expression, circulating levels and protein expression in experimental animal studies ^22, 25, 26^. MCP-1 and IL-1β deficient mice and blocking MCP-1 in rats showed impaired cerebral aneurysm formation and progression ^14, 15^. The lack of IL-1β, MCP-1, and the inhibition of MCP-1 receptor CCR2 decreased atherosclerotic formation ^16–18^. From these findings, it can be postulated that colchicine inhibiting SASP factors (Figure 6)^5, 22, 24–26, 30^ can alleviate the formation and progression of atherosclerosis and CAs ^14–18^.

MMPs released by senescent cells contribute to cardiovascular diseases via remodeling vascular tissue by degenerating extracellular matrix ^10, 47^. In our study, colchicine inhibited mRNA and protein expression of MMP-2 (Figure 6). Human CAs stained extensively for MMP-2 and MMP-9 while the circle of Willis arteries showed minimal or negative staining ^56^.

The protein expression of MMP-2 and MMP-9 was significantly upregulated in human CAs than in control arteries ^56^. The mRNA expression of MMP-2 increased with the progression of CAs and the mRNA expression of MMP-9 was up-regulated after 3 months of CAs induction in rats ^13^. Inhibiting MMP-2 and MMP-9 reduced the progression of CAs in rats ^13^. MMP-2 deficiency significantly reduced atherosclerotic plaque lesion formation in mice ^12^. MMPs can also modulate inflammation through their protease activity by post-translational processing of inflammatory cytokines such as TNF-α and IL1-β, chemokines like MCP-1 and cell adhesion molecules namely ICAM-1 ^11^. Colchicine reduced mRNA and protein expression of MMP2 *in vitro* ^5^, mitigated mRNA expression of MMP-3, MMP-9, and MMP-10 in aortas of mice ^22^ and abrogated the relative mRNA expression of MMP-9 in infarct area of myocardium after myocardial infarction in mice ^26^. These findings advocate that colchicine by inhibiting MMP-2 mRNA and protein expression can reduce tissue remodulation, and thus can mitigate the formation and progression of atherosclerotic lesions and CAs (Figure 6) ^12, 13^.

Clinical and animal experimental studies have reported the beneficial effects of colchicine on cardiovascular diseases. Colchicine reduced atherosclerotic plaque size in mice ^22^ and rabbits ^23^, attenuated aortic outward vascular remodeling in rabbits ^23^, reduced medial fibrosis in rabbits ^23^, ameliorated plaque vulnerability index in rabbits ^23^ and inhibited Carrageenan-induced thrombosis in mice ^24^. Colchicine ameliorated cardiac damage in doxorubicin-induced mice, improved cardiac structural remodeling, and attenuated mechanical dysfunction during dilated cardiomyopathy progression ^25^. Colchicine improved survival rate and inhibited heart failure after myocardial infarction in mice ^26^. Furthermore, recent clinical trials have reported that a daily dose of 0.5 mg of colchicine lowered the risk of cardiovascular events in patients with a recent myocardial infarction ^27^ and in patients with chronic coronary disease ^28^.

The observations of the current study together with previously reported findings suggest that colchicine protects against cardiovascular diseases by inhibiting cellular senescence and SASP by blocking the activation of NF-κB and MAPKs through reducing oxidative stress and subsequent DNA damage.

## Conclusion

Endothelial cell senescence contributes to the progression of cardiovascular diseases. In the current study, we show that oxidative stress activates NF-kB, MAPKs, and mTOR pathways and induces premature senescence and SASP in endothelial cells. Colchicine suppressed oxidative stress-induced senescence and SASP in HUVECs by blocking the activation of the NF-kB and MAPKs pathways.

## Limitations

Our study has some limitations. The study was performed using *in vitro* model of primary endothelial cells. In the study, we used endothelial cells from the human umbilical vein and not from the artery. Therefore, the results should be interpreted carefully.

## Author’s contributions

Conceptualization, Dilaware Khan and Sajjad Muhammad; Investigation, Huakang Zhou and Dilaware Khan; Methodology, Huakang Zhou and Dilaware Khan; Project administration, Dilaware Khan and Sajjad Muhammad; Resources, Sajjad Muhammad; Supervision, Dilaware Khan and Sajjad Muhammad; Writing – original draft, Huakang Zhou and Dilaware Khan; Writing – review & editing, Sajid Hussain, Norbert Gerdes, Carsten Hagenbeck, Majeed Rana, Jan Cornelius and Sajjad Muhammad.

## Acknowledgment

Research commission grant of Heinrich Heine University Düsseldorf 2020 and 2023 to SM EANS research funds for the vascular category 2019 and 2020 to SM Peek and Cloppenburg Stifftung 2021 to SM

## Conflict of Interest

The authors declare no conflict of interest.

## Compliance with ethics guidelines

This article does not contain any studies with human or animal subjects.

## Data Availability Statement

The data is contained within the article.

## Highlights

- Colchicine inhibited oxidative stress and oxidative stress induced DNA damage
- Colchicine prevented oxidative stress induced endothelial cell senescence
- Colchicine inhibited activation of NF-κB and MAPKs
- Colchicine mitigated mRNA expression of inflammatory markers and MMPs

## Notes

### Competing Interest Statement

The authors have declared no competing interest.

